# Pregnant Dairy Heifers Express Influenza A Virus Receptors in the Mammary Gland

**DOI:** 10.1101/2025.05.08.652757

**Authors:** Pari H. Baker, Leandre M. Glendenning, Mauricio X.S. Oliveira, Brian A. Cobb, Benjamin D. Enger, Stephanie N. Langel

## Abstract

Highly pathogenic avian influenza (HPAI) A(H5N1) virus emerged in lactating dairy cattle in March 2024, causing mastitis-related disease and infections in other farm animals and workers. Recent work identified α2,6 and α2,3-linked sialic acids (SA), which serve as influenza virus receptors, in the lactating bovine mammary gland; however, their distribution across stages of mammary growth and development remains unknown. We compared the distribution of tissue sialylation in mammary glands of prepubertal dairy calves, pregnant dairy heifers, and lactating cows. Mammary glands at all physiological stages expressed both α2,6 SA, the preferred receptor linkage for human influenza viruses, and α2,3 SA, the preferred receptor linkage for avian influenza viruses. Importantly, mammary glands of pregnant dairy heifers exhibited the highest overall expression of α2,3 SA, observed in both tissue and alveolar lumens. Our results suggest that pregnant dairy heifers, like lactating dairy cows, are susceptible to H5N1 infection in the mammary gland.

## INTRODUCTION

The emergence of highly pathogenic avian influenza (HPAI) A(H5N1) clade 2.3.4.4b virus in dairy cattle in the United States has raised significant concerns about potential cross-species transmission and the economic impact on the dairy industry [1, 2]. Lactating dairy cows have been primarily affected, with initial cases identified after reports of reduced milk yields on farms. Affected animals exhibit nonspecific clinical signs, including decreased rumination, abnormal milk characteristics, and mild respiratory distress [3-5]. In recent studies, milk from infected cows contains the highest levels of viral RNA and infectious virus compared to other samples and tissue sites [3-5]. As of May 2025, HPAI H5N1 virus has been documented in dairy cattle across 17 states, affecting at least 1,048 dairy herds [6]. The American Association of Bovine Practitioners has estimated the economic impact of H5N1 virus on dairy farms to be between $100 and $200 per clinically diseased cow, which is significant for large-scale dairy operations [7]. Additionally, this estimate does not include the long-term effects of the disease, such as premature culling [7]. The H5N1 virus outbreak on dairy farms has also led to zoonotic transmission to farm workers and other farm animals, increasing the economic and health consequences of such outbreaks.

Influenza A virus (IAV) has been rarely documented in cattle, but previous studies have observed an association between respiratory disease, increased influenza A antibody titers in serum, and reduced milk yield [8, 9]. The attachment of IAV strains to host cells require binding to terminal sialic acids (SA) of oligosaccharides found on cell surface glycoproteins [10, 11]. Sialic acids exist in various forms, and the binding specificity of IAV for different SA linkages is considered a determinant of host range, tissue tropism, and virulence [10]. For example, IAV isolated from humans and swine preferentially bind to SAs linked to galactose (Gal) by an α2,6 linkage (α2,6 SA) [12-15] and are predominant in the upper respiratory tract [10], while avian influenza virus strains preferentially bind SAs attached to Gal via an α2,3 linkage (α2,3 SA) [10, 14-17] which are commonly found in the intestinal tract of birds and the lower respiratory tract of humans [10]. Additionally, avian influenza viruses isolated from chickens and ducks differ in their affinity for the saccharide underlying the Gal residue linked to terminal α2,3 SA. Gambaryan et al. (2003) demonstrated that chicken-derived influenza viruses preferentially bind to SA-α2,3-Gal-β1,4-N-acetylglucosamine (GlcNAc), while those derived from waterfowl preferred SA-α2,3-Gal-β1,3)-N-acetylgalactosamine (GalNAc) [16]. Recent studies have determined that dairy cows express both α2,6 SA and α2,3 SA in the respiratory tract and mammary tissues [18-21]. Notably, α2,3 SA is most abundant in mammary tissue, whereas lower levels, or none at all, are detected in the trachea, bronchi, and lung alveoli [20, 21]. The expression of α2,6 SA and α2,3 SA in bovine tissue underscores the adaptability of IAVs to nontraditional species and highlights their potential for cross-species transmission [22].

The current H5N1 outbreak involving the mammary glands of lactating dairy cows raises questions about bovine mammary gland susceptibility across the dairy cow lifespan. A key outstanding question is whether non-lactating dairy heifers are susceptible to intramammary H5N1 infection, potentially serving as reservoirs by becoming infected prior to entering the lactating herd. The potential for non-lactating dairy heifers to develop intramammary H5N1 infection is particularly relevant given that they are typically housed in open or semi-open environments, increasing their exposure to wild birds. Additionally, upon entering the milking parlor after calving, actively infected heifers could introduce H5N1 into a previous H5N1 negative lactating herd. This study aimed to evaluate the sialylation of bovine mammary glands across different physiological states to identify potential periods of susceptibility to H5N1 infection. Knowledge of SA distribution in the mammary gland and the timing of susceptibility to H5N1 clade 2.3.4.4b virus infection may enable producers to implement targeted preventative measures, reduce the risk of infection within their herds, and potentially initiate treatment before the onset of clinical signs. Such efforts could help reduce environmental levels of H5N1 virus and limit zoonotic transmission in both animals and humans.

## MATERIALS AND METHODS

### Study Design

Archived mammary tissues collected from healthy mammary glands of prepubertal dairy heifer calves (n = 4; [23]), 6.5-month (n = 4) and 8.5-month (n = 4) pregnant dairy heifers (Oliveria and Enger, unpublished; Institutional Animal Care and Use Committed Protocol 2020A00000024), and lactating cows (n = 4; [24] and Montgomery and Enger, unpublished; Institutional Animal Care and Use Committed Protocol 2022A00000020) were used and examined. Mammary tissues were collected above the gland cistern, near the center of the parenchymal mass, and fixed in 10% formalin for 48 h before being embedded in paraffin.

### H&E Staining

A hematoxylin and eosin slide set was prepared to compare the morphological differences between the mammary glands of different physiological states. Paraffin embedded tissues were sectioned at 4-μm thick using a rotary microtome (model HM 340 E; Microm International GmbH), relaxed in a 42 °C water bath, and mounted to SuperfrostTM Plus microscope slides (Thermo Fisher Scientific). Slides were dried on a slide warmer for 24 hours. Dried sections were then stained with hematoxylin and eosin as previously described [25, 26]. Hematoxylin and eosin–stained sections were visualized and imaged with an Olympus BX43 microscope (Olympus Corp.) fitted with a SC180 camera (Olympus Corp) using a 4× PlanN objective.

### Lectin Histochemistry

A slide set was prepared to evaluate the degree and localization of SA expression within the mammary tissues. Briefly, tissues were sectioned and mounted as before. Tissue sections were stained with fluorophore-conjugated lectins (**Table 1**) diluted to 1 μg/ml in phosphate buffered saline and incubated overnight at room temperature. The following lectins were used to detect specific SA linkages: 1) SNA (*Sambucus nigra agglutinin*) for α2,6 SA, 2) MAL-I (*Maackia amurensis lectin I*) for SA-α2,3-Gal-β1,4-GlcNAc, and 3) MAL-II (*Maackia amurensis lectin II*) for SA-α2,3-Gal-β1,3-GalNAc (**Figure 1**). Sections were coverslipped using VECTASHIELD HardSet Antifade mounting medium (Vector Laboratories, Newark, CA), and imaged using a Leica SP5 Laser-Scanning Confocal Microscope using a 10× objective. Images were analyzed using ImageJ Software to quantify the mean fluorescence intensity (MFI) of each fluorophore-conjugated lectin. Each dot on the lectin quantitation graphs represents the average MFI of six images per tissue section (n=4 animals/group). Data expressed as the binding of each lectin as a ratio of Concanavalin A (ConA), which recognizes α-linked mannose and acts as an internal N-glycosylation positive control.

**Table 1.**
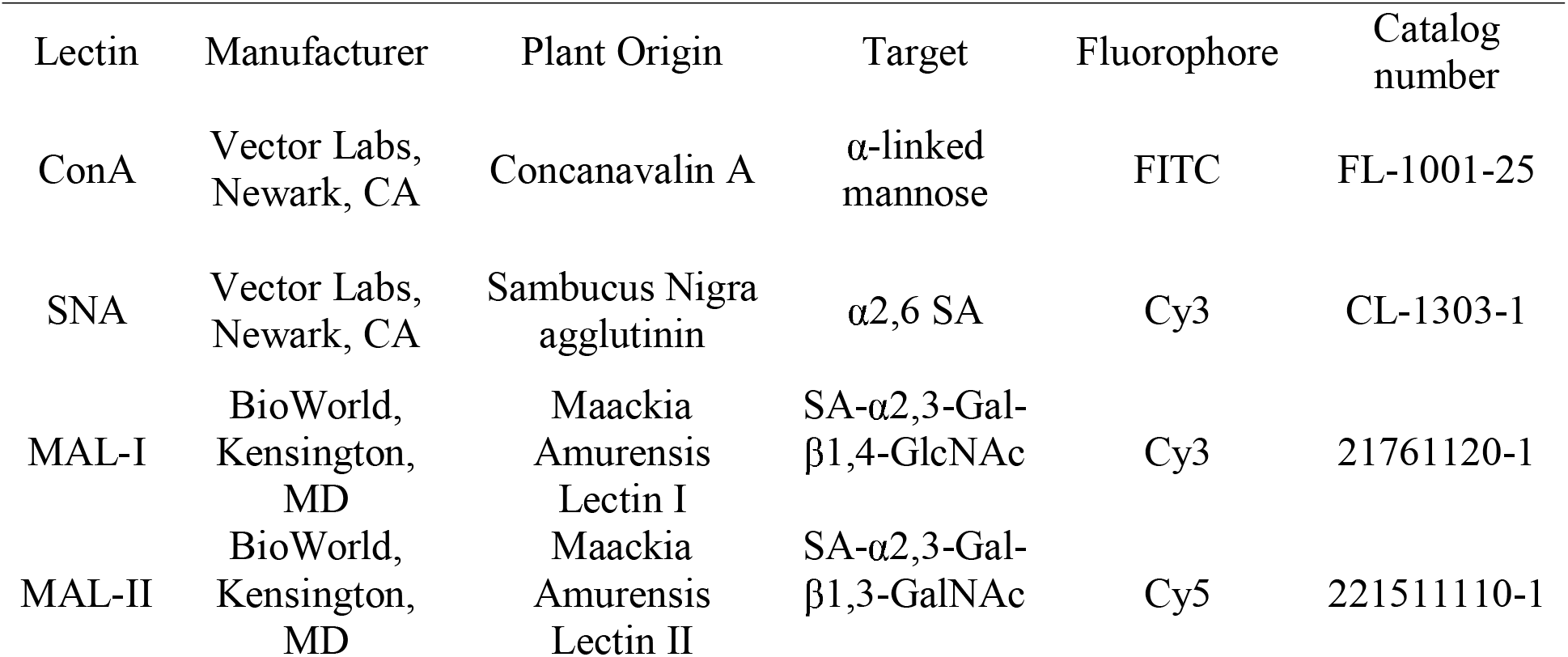
Lectins, manufacturer, plant origin, target sialic acid, fluorophore and catalog number used to label sialic acids in mammary tissue.

| Lectin | Manufacturer | Plant Origin | Target | Fluorophore | Catalog number |
| --- | --- | --- | --- | --- | --- |
| ConA | Vector Labs, Newark, CA | Concanavalin A | α-linked mannose | FITC | FL-1001-25 |
| SNA | Vector Labs, Newark, CA | Sambucus Nigra agglutinin | α2,6 SA | Cy3 | CL-1303-1 |
| MAL-I | BioWorld, Kensington, MD | Maackia Amurensis Lectin I | SA-α2,3-Gal-β1,4-GlcNAc | Cy3 | 21761120-1 |
| MAL-II | BioWorld, Kensington, MD | Maackia Amurensis Lectin II | SA-α2,3-Gal-β1,3-GalNAc | Cy5 | 221511110-1 |

**Figure 1.**
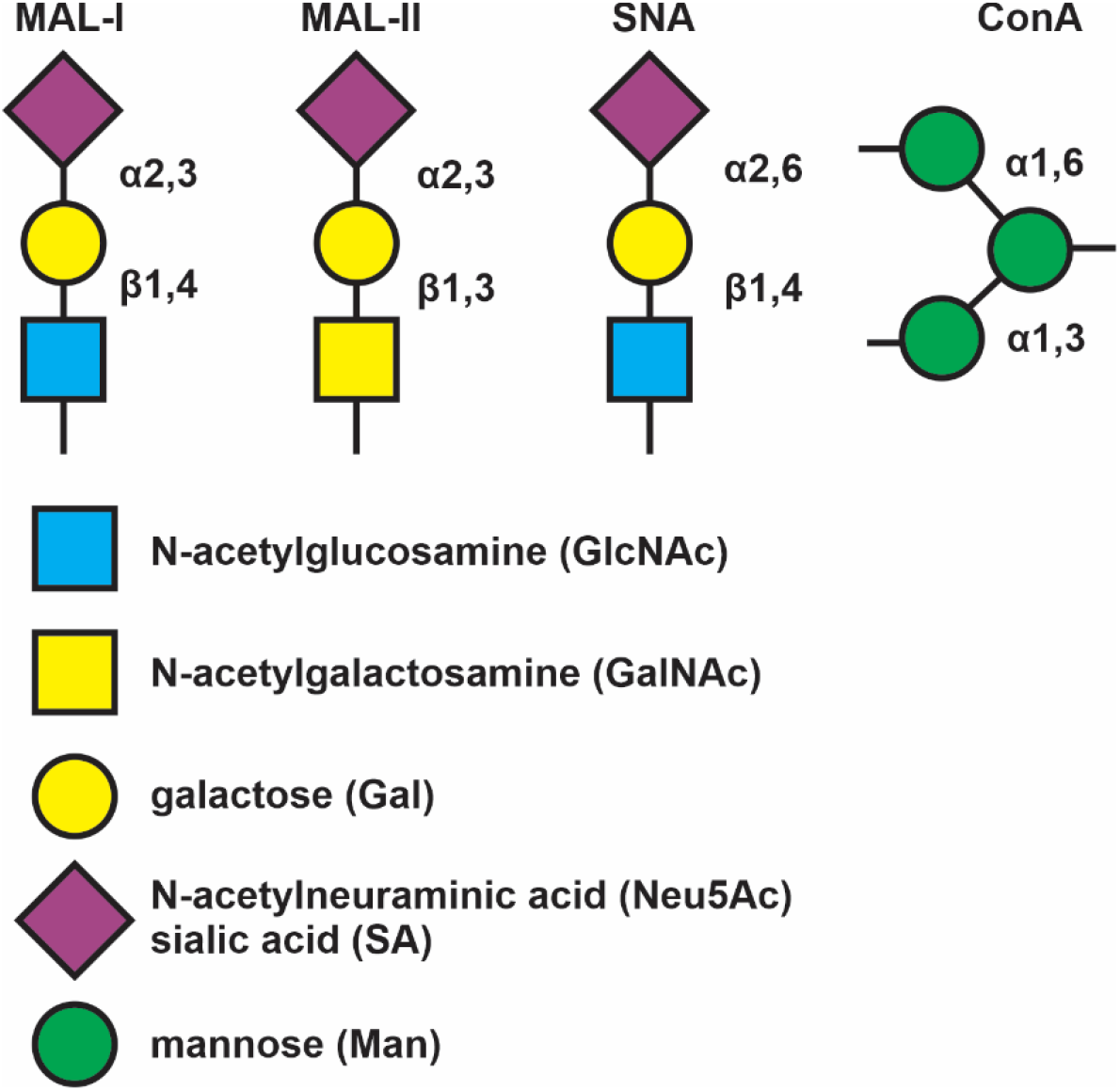
Annotation of sialic acid binding lectins. Abbreviations and symbols: Mannose (Man, green circles), sialic acid (Sia, pink diamonds), *N-*acetylglucosamine (GlcNAc, blue squares), galactose (Gal, yellow circles), *N-*acetylgalactosamine (GalNAc, yellow squares). The Symbolic Nomenclature for Glycans (SNFG) is used.

### Statistical Analyses

Statistical analyses include two-way analysis of variance (ANOVA) (mixed model) with Tukey’s multiple comparisons test. Statistical analyses were conducted on all individual data points (6 images per animal, 4 animals per group; 24 data points total per group). Statistical significance was defined as *P* ≤ 0.05. All graphs and statistical analyses were performed using GraphPad Prism software version 10.0 (GraphPad Software, San Diego, CA).

## RESULTS

### Bovine mammary gland morphology varies between physiological states

Mammary tissues of 3-month-old calves contained rudimentary epithelial structures that were often stratified. Stromal tissue occupied most of the calf mammary tissue area, containing dense connective tissue and interspersed adipocytes (**Figure 2A**). Mammary tissues of pregnant dairy heifers were markedly reorganized in preparation for lactation (**Figure 2B-C**). Stromal tissue areas were reduced while epithelial areas were increased. Additionally, luminal space was greater than in calf mammary tissues. Epithelial lobular areas and luminal space abundance increased as gestation advanced, while stromal tissue areas became reduced. Additionally, adipocytes and the dense connective tissues became less evident and nearly absent by 8.5 months of gestation (**Figure 2C**). In lactating mammary tissues, fully differentiated secretory epithelial cells were evident within well-formed alveoli containing large luminal spaces; stromal tissue areas were limited (**Figure 2D**).

**Figure 2.**
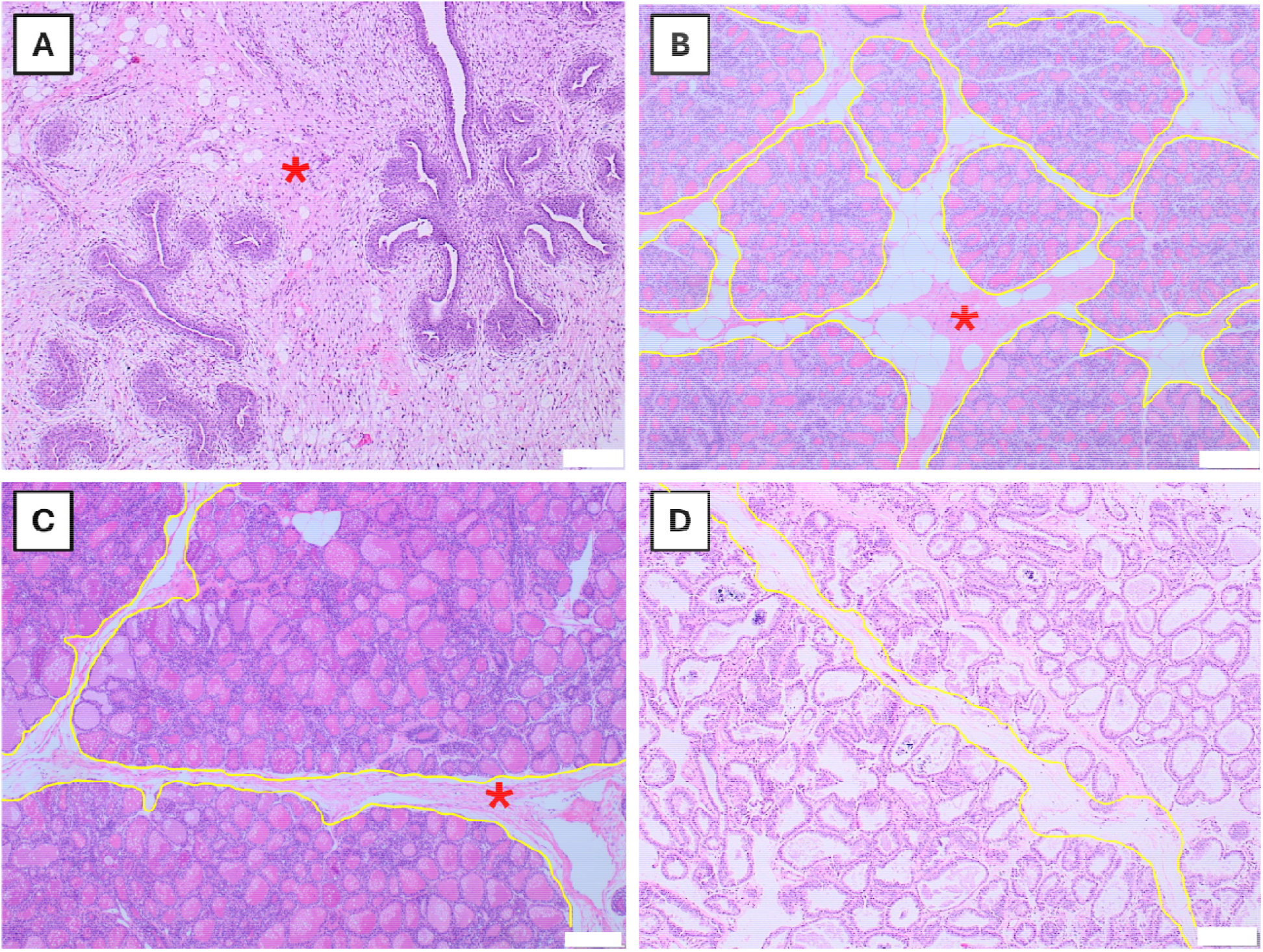
Histological evaluation of the bovine mammary gland across distinct physiological stages of growth and development. Progression of mammary growth and development can be observed from predominantly stromal tissue (red asterisk) and limited mammary epithelium seen in the calf **(A)**, 6.5-mo pregnant animals exhibiting more adipose tissue areas and less lobular area (yellow outline) compared to 8.5-mo pregnant animals **(B and C, respectively)**, and the lactating mammary tissue exhibiting minimal adipose tissue, large lobular area, well-developed alveoli with large luminal areas **(C)**. Scale bar = 200 µm.

### Lactating mammary tissue exhibited the highest levels of α2,6 SA expression

We evaluated the distribution of α2,6 SA (SNA) and α2,3 SA glycoforms, specifically the SA-α2,3-Gal-β1,4-GlcNAc common in complex N-glycans (MAL-I), and SA-α2,3-Gal-β1,3-GalNAc common in O-linked glycans (MAL-II), in bovine mammary tissues across distinct physiological states using lectin-based fluorescent histochemistry as indicated. We found that mammary tissue from lactating cows had significantly higher expression of α2,6 SA compared to mammary tissue from prepubescent calves (*P* < 0.01), early gestation (6.5-month pregnant) heifers (*P* < 0.01) and late gestation (8.5-month pregnant) heifers (*P =* 0.004); **Figure 3A**. In prepubertal calf mammary tissue, α2,6 SA expression was polarized to the apical side of the epithelial structures with minor to moderate expression in the intralobular stroma, particularly being associated with endothelium (**Figure 3B**). Expression of α2,6 SA in mammary tissue of pregnant heifers was polarized to the apical side of epithelial cells, being highly abundant within alveolar lumens, and was faintly observed in endothelial structures (**Figure 3C-D**). In the lactating cow mammary tissue, α2,6 SA was polarized to the apical membrane of the mammary epithelium and remained evident in the endothelium (**Figure 3E**).

**Figure 3.**
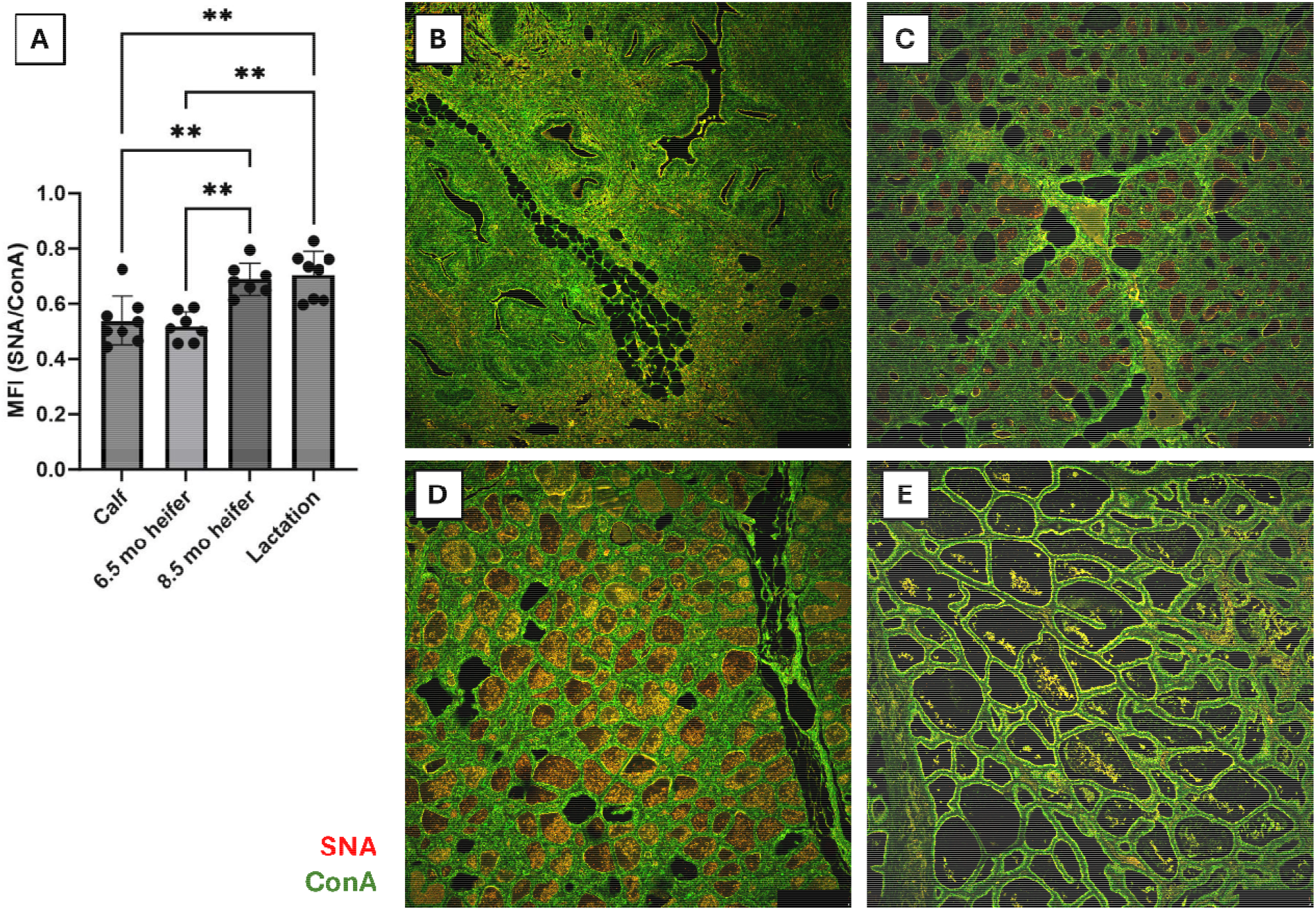
Expression of α2,6 SA within mammary tissue at different stages of development. Data expressed as the mean fluorescence intensity (MFI) of SNA normalized to the MFI of ConA **(A)**. Each dot on the lectin quantitation graph represents the average MFI of six images per tissue section (n=4 animals/group). A two-way Analysis of Variance (ANOVA) was used to compare treatment groups, and error bars represent the standard deviation of the mean MFI for each group. Mammary gland tissues from **(B)** prepubescent dairy calves, **(C)** 6.5-month pregnant, **(D)** 8.5-month pregnant heifers and **(E)** lactating dairy cows were stained with lectins Cy3-labeled SNA (red) and FITC-labeled ConA (green) and incubated overnight. Color overlay to illustrate colocalization (yellow). SNA, *Sambucus nigra* lectin; ConA, Concanavalin A. Scale bar = 250 µm. Asterisks indicate statistically significant differences (\**P* ≤ 0.05, \*\**P* ≤ 0.01, \*\*\**P* ≤ 0.001, \*\*\*\**P* ≤ 0.0001).

### Bovine mammary tissue at distinct physiological states had minimal expression of SA-α2,3Gal-β1,4-GlcNAc N-glycans

There was no significant difference in SA-α2,3-Gal-β1,4-GlcNAc expression between bovine mammary tissue at distinct physiological states (**Figure 4A**). Expression of SA-α2,3-Gal-β1,4-GlcNAc, detected using MAL-I, was virtually absent across all mammary tissues (**Figure 4C-F**). In addition, the relative signal of MAL-I when normalized to ConA was significantly lower than that of both SNA/ConA (*P* < 0.01) and MAL-II/ConA (*P* < 0.01); **Supplemental Figure S1**.

**Figure 4.**
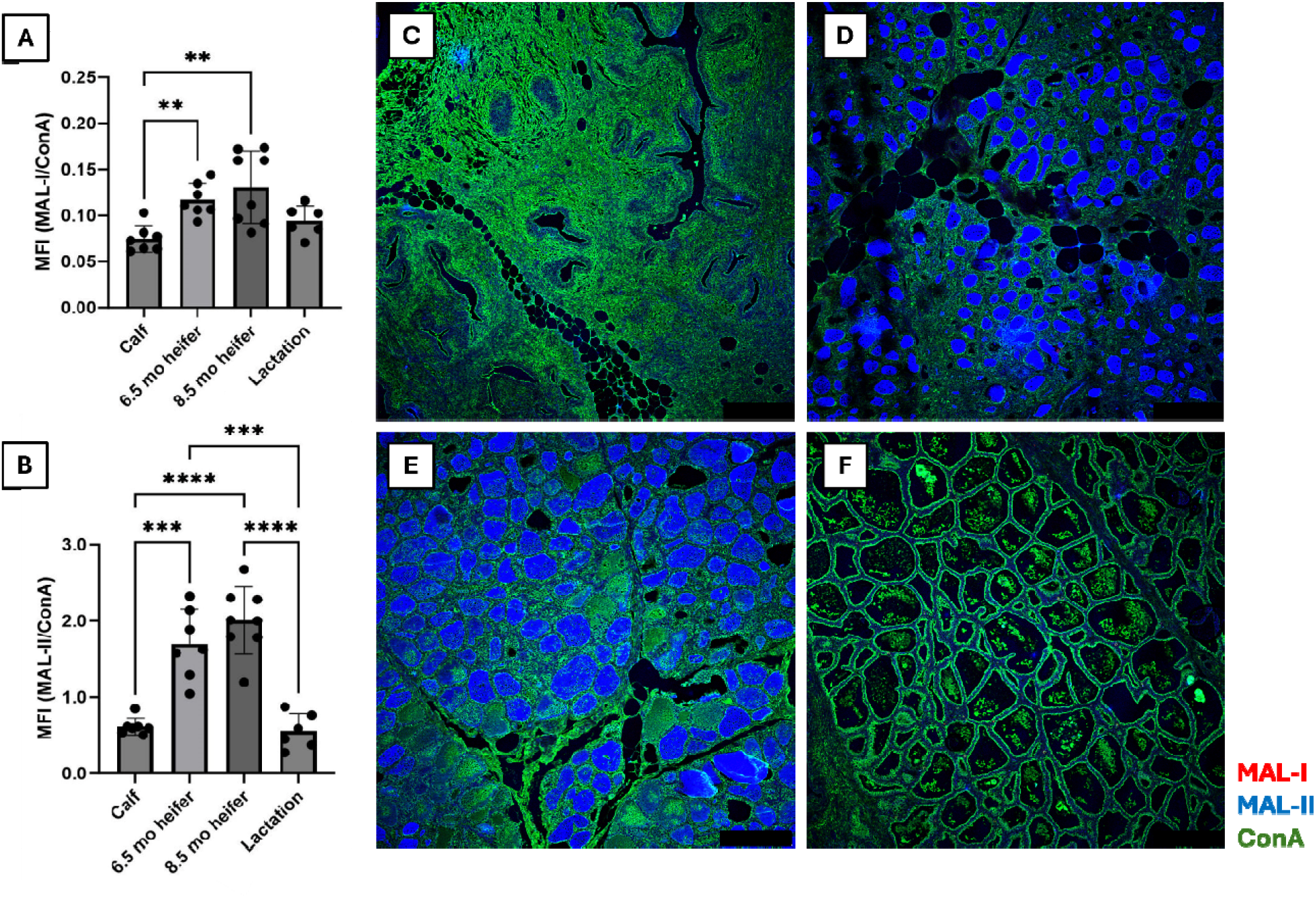
Expression of α2,3 SA within mammary tissue at different stages of development. Data expressed as the mean fluorescence intensity (MFI) of MAL-I normalized to the MFI of ConA **(A)**, and the MFI of MAL-II normalized to the MFI of ConA **(B)**. Each dot on the lectin quantitation graph represents the average MFI of six images per tissue section (n=4 animals/group). A two-way Analysis of Variance (ANOVA) was used to compare treatment groups, and error bars represent the standard deviation of the mean MFI for each group. Mammary gland tissues from **(C)** prepubescent dairy calves, **(D)** 6.5-month pregnant heifers, **(E)** 8.5-month pregnant heifers and **(F)** lactating dairy cows were stained with lectins Cy3-labeled MAL-I (red), Cy5-labeled MAL-II (blue), and FITC-labeled ConA (green) and incubated overnight. Color overlay to illustrate colocalization with Cy-3-labeled MAL-I virtually absent in tissue. MAL, *Maackia amurensis* lectin; ConA, Concanavalin A. Scale bar = 250 µm.

### Mammary glands from pregnant heifers expressed the greatest levels of SA-α2,3-Gal-β1,3-GalNAc O-glycans

We observed that the abundance of SA-α2,3-Gal-β1,3-GalNAc, detected using MAL-II, was significantly greater in mammary glands of 6.5-month and 8.5-month pregnant dairy heifers compared to prepubescent calves (*P* ≤ 0.01) and lactating cows (*P* ≤ 0.01; **Figure 4B**). The location of SA-α2,3-Gal-β1,3-GalNAc expression in prepubertal dairy calf mammary tissue was diffusely present in the rudimentary epithelial structures (**Figure 4C**), while SA-α2,3-Gal-β1,3-GalNAc was heavily concentrated within the alveolar lumens of the 6.5-and 8.5-month pregnant heifer mammary tissues; minor to moderate expression in endothelial structures and intralobular stroma was also observed **(Figure 4D-E**). In contrast, SA-α2,3-Gal-β1,3-GalNAc expression in lactating tissue was most apparent in intralobular stroma areas between the alveoli, with minimal presence in the alveolar lumens (**Figure 4F**).

## DISCUSSION

Our central objective was to evaluate the distribution of α2,6 and α2,3 SA in bovine mammary glands at different physiological stages. Given the detection of HPAI H5N1 virus in the mammary glands and milk of lactating cows, evaluating SA distribution in the mammary glands during other physiological states is warranted. Recent studies have reported that mammary tissue from lactating dairy cows expresses both α2,6 and α2,3 SA [18-20]. Kristensen et al. (2024) reported the presence of α2,6 SA and SA-α2,3-Gal-β1,3-GalNAc, the preferred SA form for influenza viruses in ducks [16], in the lactating mammary gland, with widespread distribution in the alveoli and more limited expression in the ducts. In contrast, no staining was observed for SA-α2,3-Gal-β1,4-GlcNAc, the SA form preferentially bound by influenza viruses isolated from chickens [16]. The complementary work by Nelli et al. (2024) revealed that uninfected lactating mammary tissue expressed SA-α2,3-Gal-β1,3-GalNAc in the secretory alveoli, with localization polarized to the apical side of the MECs, while SA-α2,3-Gal-β1,4-GlcNAc was minimally detected [20]. Additionally, α2,6 SA expression was predominately located to the apical side of the mammary epithelium. Unlike the observations from Kristensen et al. (2024), Nelli et al. observed abundant α2,6 SA expression in the epithelial cells lining the interlobular ducts [20]. However, a subsequent study by Carrasco et al. (2024) observed an absence of α2,6 SA expression and minimal HA binding from the human influenza strain (A/Puerto Rico/8/34 H1N1) [21], which contradicts both our findings and those of previous studies [19, 20]. Despite these varying results, the collective findings highlight the dynamic nature of glycan profiles in the bovine tissues and underscore the need for in vivo studies to define mammary gland susceptibility to influenza viruses.

We found that mammary tissue from lactating cows had significantly greater α2,6 SA abundance than to mammary tissue of prepubescent calves and 6.5-and 8.5-month pregnant heifers. Localization of SAs with a α2,6 linkage in mammary tissue across all physiological states appeared polarized to the apical side of the MECs, an observation that coincides with those from previous studies [19, 20]. Interestingly, we also noted the presence of α2,6 SA in the intralobular stroma, underlying the endothelial cells or lymphatics. However, further characterization of the stromal cellular components is needed to confirm these observations and investigations into the significance of SA presence in endothelium is warranted.

Coinciding with previous studies [18-20], we found that mammary tissue at each physiological state had minimal SA-α2,3-Gal-β1,4-GlcNAc glycoforms. Within all mammary tissue, SA-α2,3-Gal-β1,3-GalNAc (MAL-II) predominated, followed by α2,6 SA (SNA) and SA-α2,3-Gal-β1,4-GlcNAc (MAL-I). When comparing the two α2,3 SA glycoforms, the SA-α2,3-Gal-β1,3-GalNAc, was significantly greater relative to ConA, compared to SA-α2,3-Gal-β1,4-GlcNAc. The lower abundance of SA-α2,3-Gal-β1,4-GlcNAc, the receptor for chicken-derived influenza viruses, compared to SA-α2,3-Gal-β1,3-GalNAc, the receptor for duck-derived influenza viruses, may help explain why the currently circulating HPAI H5N1 clade 2.3.4.4b virus, which originated in waterfowl, has emerged in lactating dairy cattle [27]. Previous studies have reported that avian influenza viruses adapted to ducks are less likely to infect chickens under experimental conditions, and conversely, avian influenza viruses infecting chickens are less likely to infect ducks [17, 28, 29]. The presence or absence of specific α2,3 SA glycoforms in the mammary gland could provide insight into the mechanisms of interspecies transmission of avian influenza viruses from ducks or chickens, as well as the tissue-specific susceptibility of mammalian hosts.

Additionally, we found that 6.5-and 8.5-month pregnant dairy heifers had significantly greater abundance of SA-α2,3-Gal-β1,3-GalNAc in their mammary gland compared to prepubescent calves and lactating cows. Expression of SA-α2,3-Gal-β1,3-GalNAc was highly concentrated in the alveolar lumens of the mammary tissue from 6.5-and 8.5-month pregnant dairy heifers, with weak cytoplasmic staining observed, suggesting diffuse distribution throughout the mammary epithelium. Mild to moderate expression of SA-α2,3-Gal-β1,3-GalNAc was noted in the intralobular stroma of pregnant mammary tissue; however, it appeared markedly pronounced in the stroma of the lactating mammary gland. In the lactating mammary tissue, SA-α2,3-Gal-β1,3-GalNAc had minimal expression in the alveolar lumens, an observation different than that reported by Kristensen et al. (2024) and Nelli et al. (2024) which could be due to the state of the mammary gland prior to collection, whether the mammary gland was full of milk or devoid of milk at the time of collection [19, 20]. MAL-II staining within the intralobular stroma of lactating mammary glands suggests potential localization to endothelial cells and lymphatic structures, which are prominent components of the stromal compartment. These data suggest that SA expression in the mammary gland varies depending on the reproductive life stage of the animal which could influence intramammary infection risk.

It is important to note that MAL-II preferentially binds O-linked glycans due to the presence of GalNAc residues common to all mucin-type O-glycans; however, MAL-II binding alone cannot unambiguously determine whether the binding target is an N- or O*-*linked glycan [30]. Further validation is needed to determine whether staining by MAL-II is a result of the presence of N*-*linked or O-linked SA-α2,3-Gal-β1,3-GalNAc. The majority of MAL-II signal was observed within the luminal compartment of the pregnant heifer mammary gland, a period during which the mammary gland undergoes critical remodeling and cellular differentiation in preparation for lactation [31]. This is an important consideration as bovine colostrum contains mucin [32], a large glycoprotein rich in O-linked glycans [33]. Future work is needed to understand whether colostrum from primiparous dairy heifer specifically contains O-linked glycans. Regardless, MAL-II signal was still detected in the intralobular stroma of mammary tissue from pregnant heifers, raising concerns about their susceptibility of dairy heifers to intramammary H5N1 virus infection. Further investigation is warranted to determine whether the developing mammary gland in primigravid heifers is susceptible to intramammary H5N1 infection and mastitis, resulting in reduction of future milk production, and increased risk of viral transmission within the dairy herd.

## CONCLUSION

Overall, our data indicates that SA expression in the bovine dairy mammary gland varies across different physiological states. Further studies are warranted to determine whether expression of α2,3 SA in the pregnant dairy heifer mammary gland results in H5N1 infection and mastitis after intramammary inoculation. Additionally, assessing whether pregnant dairy heifers can serve as reservoirs for H5N1on the dairy farm should be a research priority, given their potential role in viral transmission. Preventing and mitigating such infections at the herd level could not only improve udder health and lactating performance but also serve as a critical strategy for reducing zoonotic H5N1 transmission. These efforts are essential to safeguarding both animal and public health, while ensuring the long-term sustainability of dairy production.

## ACKNOWLEDGMENTS

This work was supported by Case Western Reserve University School of Medicine (S.N.L.) and The Ohio State University (B.D.E.).

