## Supplemental Figure 1 for "Pregnant Dairy Heifers Express Influenza A Virus Receptors in the Mammary Gland"

**Supplemental Figure S1.**

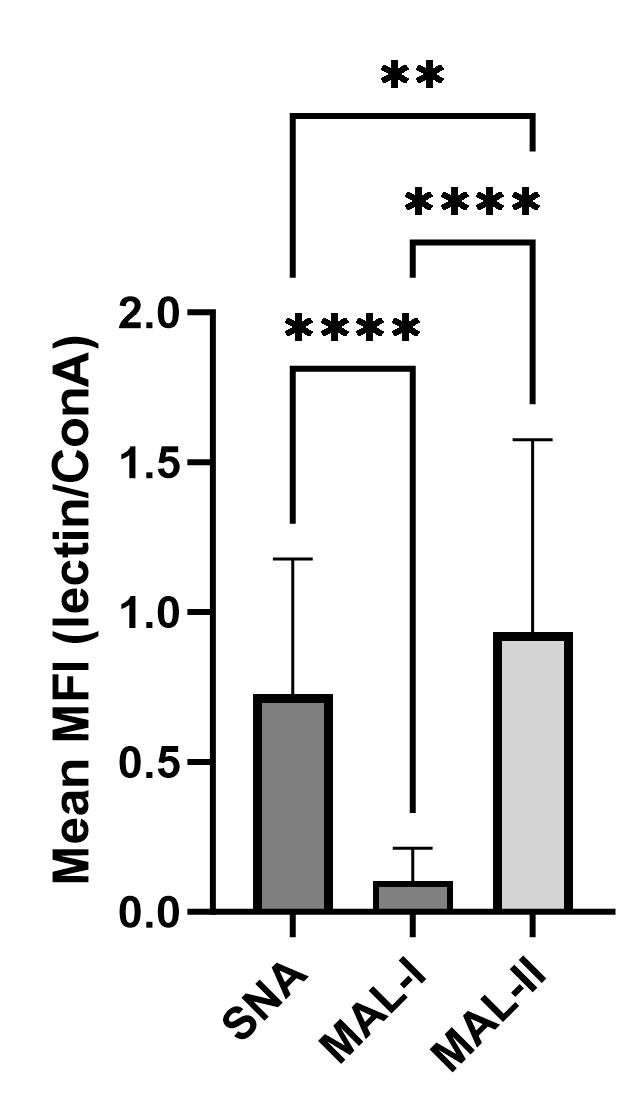

**Supplemental Figure S1. Mean MFI (mean fluorescence intensity) expression of each lectin as a ratio over ConA.** The following lectins were used for the following SA specificities: 1) SNA (*Sambucus nigra agglutinin*) for α2,6 SA, 2) MAL-I (*Maackia amurensis lectin I*) for SA-α2,3-Gal-β1,4-GlcNAc, and 3) MAL-II (*Maackia amurensis lectin II*) for SA-α2,3-Gal-β1,3-GalNAc. Asterisks indicate statistically significant differences (**P* ≤ 0.05, ***P* ≤ 0.01, ****P* ≤0.001, *****P* ≤ 0.0001).
